# Reduction of spontaneous cortical beta bursts in Parkinson’s disease is linked to symptom severity

**DOI:** 10.1101/775494

**Authors:** Mikkel C. Vinding, Panagiota Tsitsi, Josefine Waldthaler, Robert Oostenveld, Martin Ingvar, Per Svenningsson, Daniel Lundqvist

**Affiliations:** NatMEG, Department of Clinical Neuroscience, Karolinska Institutet, Stockholm, Sweden; Neuro Svenningsson, Department of Clinical Neuroscience, Karolinska Institutet, Stockholm, Sweden; Department of Neurology, University Hospital Marburg, Germany; Donders Institute for Brain, Cognition and Behaviour, Radboud University, Nijmegen, Netherlands; Section of Neuroradiology, Karolinska University Hospital, Stockholm, Sweden

**Keywords:** Parkinson’s disease, beta bursts, beta band, bradykinesia, resting-state.

## Abstract

Parkinson’s disease is characterized by a gradual loss of dopaminergic neurons, which are associated with altered neuronal activity in the beta band (13-30 Hz). Assessing beta band activity typically involves transforming the time-series to get the power of the signal in the frequency-domain. Such transformation assumes that the time-series can be reduced to a combination of steady-state sine-and cosine waves. However, recent studies have suggested that this approach masks relevant biophysical features in the beta band activity—for example, that the beta band exhibits transient bursts of high-amplitude activity.

In an exploratory study we used magnetoencephalography (MEG) to record cortical beta band activity to characterize how spontaneous cortical beta bursts manifest in Parkinson’s patients ON and OFF dopaminergic medication, and compare this to matched healthy controls. From three minutes of MEG data, we extracted the time-course of beta band activity from the sensorimotor cortex and characterized high-amplitude epochs in the signal to test if they exhibited burst like properties. We then compared the rate, duration, inter-burst interval, and peak amplitude of the high-amplitude epochs between the Parkinson’s patients and healthy controls.

Our results show that Parkinson’s patients OFF medication had a 6-17% lower beta bursts rate compared to healthy controls, while both the duration and the amplitude of the bursts were the same for Parkinson’s patients and healthy controls and medicated state of the Parkinson’s patients. These data thus support the view that beta bursts are fundamental underlying features of beta band activity, and show that changes in cortical beta band power in PD can be explained primarily by changes in the underlying burst rate. Importantly, our results also revealed a relationship between beta bursts rate and motor symptom severity in PD: a lower burst rate scaled with increased in severity of bradykinesia and postural/kinetic tremor. Beta burst rate might thus serve as neuromarker for Parkinson’s disease that can help in the assessment of symptom severity in Parkinson’s disease or evaluate treatment effectiveness.

## 1 Introduction

Parkinson’s disease is a neurodegenerative disease that, most often, initially manifests with motor symptoms such as tremor, rigidity, and bradykinesia. The neurodegenerative process is characterized by a loss of dopamine and death of dopaminergic neurons throughout the basal ganglia-thalamic-cortical system (Rodriguez-Oroz et al., 2009; Kalia and Lang, 2015). The dopamine loss leads to widespread functional changes in brain activity; for instance, throughout the basal ganglia-thalamic-cortical network, oscillatory activity in the beta band (13–30 Hz) exhibits systematic disease-related changes in Parkinson’s disease (Jenkinson and Brown, 2011). The direct influence of dopamine has for example been demonstrated to increase beta band power in the sub-thalamic nucleus (STN) when Parkinson’s patients are OFF dopaminergic medication as compared to ON medication (Alonso-Frech et al., 2006; Kühn et al., 2006; Mallet et al., 2008; Giannicola et al., 2010; Neumann et al., 2017). Increased beta power in the STN and the basal ganglia has further been linked to increased severity of bradykinesia and rigidity in Parkinson’s patients (Kühn et al., 2006; Martin et al., 2018). Disease-related changes in the beta band are found not only in STN and basal ganglia in Parkinson’s patients but is also present in the cortex, from where brain activity can be recorded non-invasively while patients are at rest, using magnetoencephalography (MEG) and electroencephalography (EEG).

Studies using MEG to assess neural activity while the participants were at rest show that Parkinson’s patients have decreased cortical beta power compared to healthy controls (Bosboom et al., 2006; Heinrichs-Graham et al., 2014). However, in the early stages of Parkinson’s disease, there seems to be an increase in beta power at rest compared to healthy controls (Pollok et al., 2012). Treatments for Parkinson’s disease also seems to be effective through modulation of the cortical beta activity. Administration of dopaminergic medication has been shown to increase the cortical beta power in Parkinson’s patients (Heinrichs-Graham et al., 2014; Melgari et al., 2014), suggesting that dopamine levels and the cortical beta power are inversely connected. Similarly, Parkinson’s patients treated with electrical deep brain stimulation (DBS) showed an increase in cortical sensorimotor beta power following DBS compared to off treatment (Airaksinen et al., 2012; Cao et al., 2017). However, other studies have reported that DBS leads to a broader suppression of 5-25 Hz power in frontal and sensorimotor cortex (Abbasi et al., 2018; Luoma et al., 2018).

It is currently unclear whether the different directions of these disparate findings are due to differences in the Parkinson’s patients (e.g., early-stage versus later-stage Parkinson’s disease) or if they are due to uncertainties in the methods used to quantify beta activity. Beta activity is traditionally assessed by analyzing the MEG/EEG data in the frequency-domain, using various forms of Fourier-transforms (e.g., wavelet-analysis) of the data. Fourier-based methods assume that the oscillatory activity in the time series can be resolved as a sum of steady-state sine and cosine waves of varying frequency. There is however converging evidence that the oscillatory activity in the beta band does not occur at a steady state, but rather consists of short transient bursts lasting only one to a few beta band cycles (Leventhal et al., 2012; Bartolo and Merchant, 2015; Feingold et al., 2015; Sherman et al., 2016). From the resulting power spectral densities (PSD) it is impossible to tell whether changes in beta band reflect a general change in the amplitude of steady-state oscillations, or if it reflects changes in the occurrence or amplitude of transient beta bursts. In all three cases, the output from the Fourier-transform will sum up to a shift in beta band power.

Several recent studies have explored the functional role of transient beta bursts in the motor cortex of healthy subjects. For instance, Shin et al. (2017) showed that the detection rate of a tactile stimulation was higher when the probability of a beta burst immediately before the stimulation was low, suggesting that the beta bursts exhibit a transient inhibitory effect on the processing of incoming sensory signals. The negative relationship between the probability of a beta burst and the detection rate of tactile stimulation has been demonstrated in both mice, monkeys, and humans (Sherman et al., 2016; Shin et al., 2017). Similarly, Little et al. (2018) showed a negative relationship between the probability of cortical beta bursts before a cued movement and reaction time in a cued reaction task, demonstrating that beta bursts have an inhibitory effect on outgoing movement initiation. Assessment of changes in beta activity in terms of transient bursts—rather than averaging in the frequency-domain—may contribute to a better understanding of what aspect of beta activity that changes in Parkinson’s disease due to disease and medication.

There is similar evidence on the functional role of transient beta bursts from research assessing beta band activity in midbrain structures. The overall power changes in the beta band in the STN can, for example, be explained as changes in the rate of high beta amplitude epochs (Tinkhauser et al., 2017a, 2018). The high-amplitude beta epochs in STN showed both increased rate and longer duration*s* when the patients were OFF dopaminergic medication as compared to ON medication. Lofredi et al. (2019) used similar measurements from STN in patients undergoing surgery to find a decrease in beta bursts in the period leading up to a movement in a cued reaction task. The relation between beta bursts and movement initiation makes beta burst a potential tool for understanding loss of control and slowing of movement in Parkinson’s disease (Tinkhauser et al., 2017b; Lofredi et al., 2019).

Analysis of beta activity at the level of beta bursts appear to be a functionally relevant approach for further understanding sensory-motor processing and may provide new insights into the function of the sensory-motor system that is lost in average based analysis method. Assessment of spontaneous beta bursts in Parkinson’s patients from non-invasive recordings, such as MEG might, therefore, provide a more sensitive assessment on how the beta band activity changes due to the disease and may help to resolve the apparently conflicting results that emerge when assuming beta band activity consist of steady-state beta oscillations.

In this study, we used non-invasive MEG measurements from Parkinson’s patients OFF and ON dopaminergic medication, and measurements from matched healthy controls, to investigate the occurrence of spontaneous transient beta bursts in the sensorimotor cortex. Our primary aim was to compare the characteristics (such as duration, amplitude, rate) of spontaneous beta burst in the sensorimotor cortex of Parkinson’s patients to healthy controls. Our secondary aim was to explore whether any of the beta bursts characteristics changed with the presence of dopaminergic medication. Finally, a third aim was to investigate whether any of the beta bursts characteristics were linked to the severity of disease symptoms in Parkinson’s disease.

## 2 Materials and methods

### 2.1 Participants

20 patients diagnosed with Parkinson’s disease (age 41–85; five female) and 20 healthy controls (age 54–76; eight female) participated in the study. The study was approved by the regional ethics committee (Etikprövningsnämden Stockholm, DNR: 2016/911-31/1) and followed the Declaration of Helsinki. All participants gave written informed consent before participating.

The patients were recruited from the Parkinson’s Outpatient Clinic, Department of Neurology, Karolinska University Hospital, Stockholm, Sweden. The inclusion criteria for the Parkinson’s patients were a diagnosis of typical Parkinson’s disease according to the United Kingdom Parkinson’s Disease Society Brain Bank Diagnostic Criteria with Hoehn and Yahr stage 1-3 (Hoehn and Yahr, 1967), under treatment with Levodopa, Catechol-O-methyltransferase inhibitor (COMT) inhibitors, Monoaminoxidase-B (MAO-B) inhibitors, or dopamine receptor agonists. Besides the diagnosis of Parkinson’s disease, the patients were healthy according to a physical examination.

Healthy controls were recruited among healthy participants who previously had participated in studies within the preceding year, or amongst the patients’ spouses.

Exclusion criteria for both groups were a diagnosis of major depression, dementia, history or presence of schizophrenia, bipolar disorder, epilepsy, or history of alcoholism or drug addiction according to the *Diagnostic and Statistical Manual of Mental Disorders DSM-V* (American Psychiatric Association, 2013). Additional exclusion criteria for the healthy controls were a diagnosis of Parkinson’s disease or any form of movement disorder.

One patient canceled the participation in the study due to severe tremor in the OFF-medication state. One healthy control was excluded from analysis due to insufficient quality of the MEG recording. The analysis includes 19 patients and 19 healthy controls.

### 2.2 Procedure

The patients were instructed to omit their morning dose of dopaminergic medication on the day of participation. Thus, the OFF state was defined as a withdrawal period of 12 hours after the last dopaminergic medication. Patients were further instructed to bring their prescribed dose of medication, which they had to take during the experiment. All patients followed the instructions.

Preparation for the MEG recordings began as soon as the participants were briefed about the procedure and signed the written informed consent. The recordings consisted of three minutes where the participants sat with their eyes closed in the MEG scanner. Text on a screen placed in front the participants initially instructed the participants to close their eyes. Participants were instructed not to open their eyes before being told to, and to avoid moving until they were allowed to open their eyes. The recordings began once the experimenter through video observation had assured that participant’s eyes were closed. The participants then did two unrelated tasks in the same recording session consisting of an active tapping task (Vinding et al., *in prep*.) and a task with passive movements (Vinding et al., 2019). Each MEG recording session took about one hour.

When the first session was over, participants had a break outside the scanner. During the break, the participants performed the neurological tests described below, and the patients took medication. The second MEG measurement began approximately one hour after medication. The healthy controls did not take any medication but had a similar duration break and measured twice to accommodate the potential effect of the fixed order of the OFF-ON measurements in patients.

Motor function was assessed in all participants using the motor subscale of the Movement Disorder Society’s Unified Parkinson’s Disease Rating Scale (MDS-UPDRS-III) (Goetz et al., 2007), by neurologists certified in the use of MDS-UPDRS. Patients were assessed immediately after the first MEG session in the OFF state and again after the second MEG session ON medication. Montreal Cognitive Assessment (MoCA) test was done in the ON state.

### 2.3 MEG recordings

MEG data were recorded with an Elekta Neuromag TRIUX 306-channel MEG system, with 102 magnetometers and 102 pairs of orthogonal planar gradiometers, inside a two-layer magnetically shielded room (model Ak3B, Vacuumschmelze GmbH), with internal active shielding active to suppress electromagnetic artifacts. Data were recorded at 1000 Hz with an online 0.1 Hz high-pass filter and 330 Hz low-pass filter. The subjects’ positions and movements inside the MEG scanner were measured during recordings with head-position indicator coils attached to subjects’ heads. The location of the coils—and additional points giving a representation of the subjects’ head shape—was digitalized with a Polhemus Fastrak motion tracker before the measurements. The head shapes were later used to co-register MEG data and structural MRI. Horizontal and vertical electrooculogram (EOG) and electrocardiogram (ECG) were recorded simultaneously with the MEG.

### 2.4 Data processing

MEG data were processed off-line by applying temporal signal space separation (tSSS) to suppress artifacts from outside the scanner helmet and correct for head movement during the recordings (Taulu and Simola, 2006). The tSSS had a buffer length of 10 s and a cut-off correlation coefficient of 0.95. Movement correction was done by shifting the head position to a position based on the median head position during the recording. We then did an independent component analysis (ICA) for each subject using the *fastica* algorithm (Hyvarinen, 1999) implemented in MNE-Python (Gramfort et al., 2013) in Python 2.7. Components related to saccadic eye-movements and heartbeats were identified based on their correlation with the EOG or ECG and removed from the data.

We then applied source reconstruction to the data using noise weighted minimum-norm estimates (dSPM) (Dale et al., 2000). The noise covariance matrix was estimated from two minutes of empty room data recorded before each session. The source space consisted of 5124 evenly spaced points sampled across the white matter surfaces. The surfaces were obtained with the automatic routine for extracting cortical surfaces in Freesurfer (Dale et al., 1999) from individual T1 weighted MRI that were obtained on a GE Discovery 3.0 T or a Siemens Prisma 3.0 T MR scanner. One subject did not complete an MR scan, so we used an MRI template (Holmes et al., 1998) warped to the subject’s head shape as a substitute. From the MRI, we obtained the inner skull boundary, which was used to create a single compartment volume conductor model to estimate the forward model.

The cortical surface was then segmented into anatomical labels based on the automatic labeling algorithm in Freesurfer (Destrieux et al., 2010). Based on the labels, we extracted data from all point within a region of interest (ROI) consisting of the pre-and post-central gyri and central sulcus of the left hemisphere (Fig. 1). We then obtained a combined ROI time course as the first right-singular vector of a singular value decomposition of the source time courses within the ROI, with the sign of the vector normalized relative to the source orientations.

**Figure 1:**
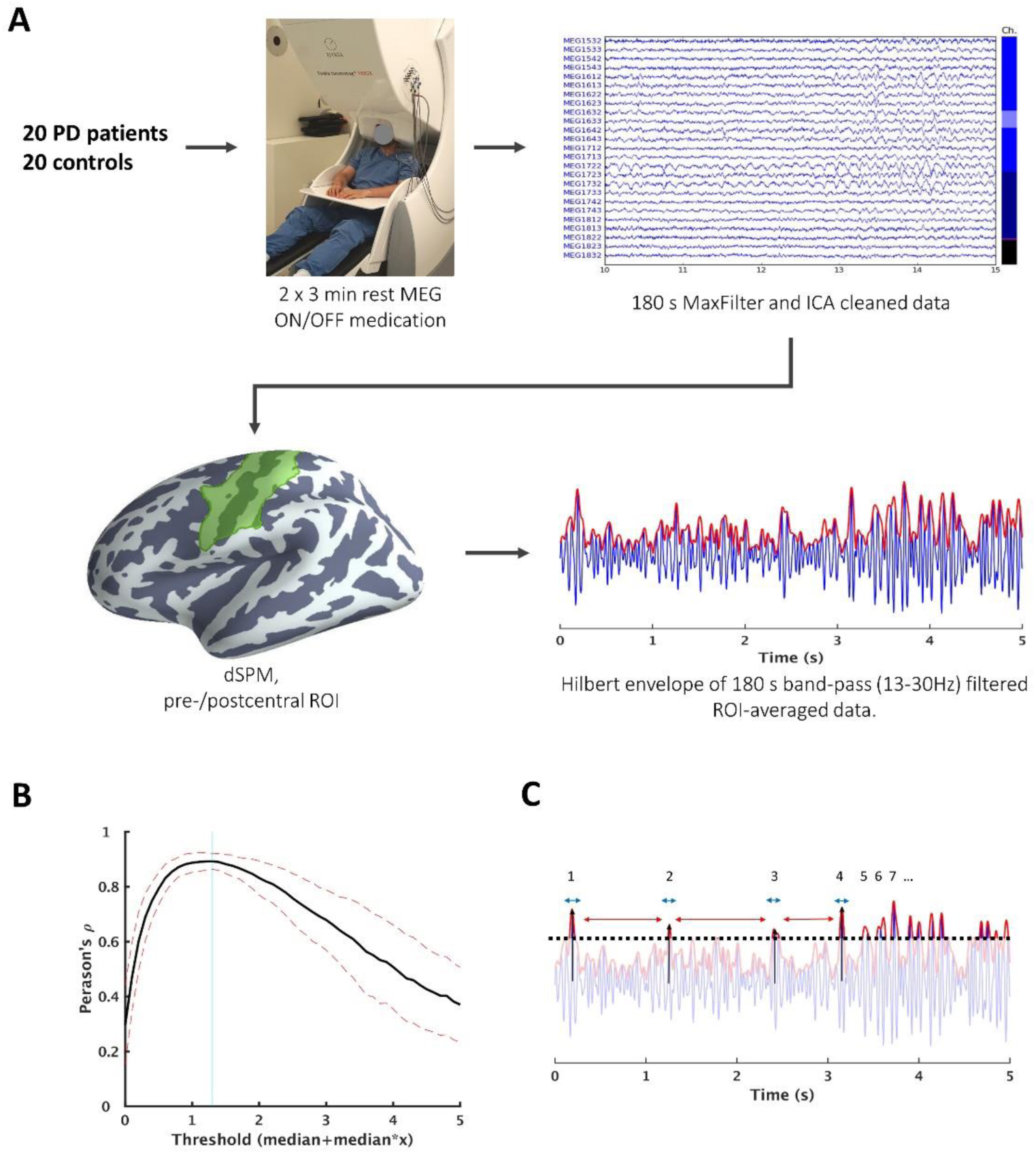
Overview of data processing from raw MEG data to characterizing beta bursts. A) We recorded three minutes of resting-state MEG. Raw MEG data were first processed with tSSS and ICA to remove artifacts. We then did a dSPM source reconstruction and extracted the time-series from an ROI consisting of the pre-/postcentral gyri and central sulcus. The ROI time-series was filtered to the beta range (13-30 Hz) and Hilbert-transformed. B) High-amplitude epochs were determined based on a threshold defined as the cutoff that had the highest correlation between the number of epochs and amplitude in consecutive 3.0 s segments. The vertical line indicates the threshold used in the analysis. C) Once the threshold was defined, we compared four features of the high-amplitude epochs: *rate* (i.e., count occurrence high-amplitude epochs), *duration* (blue arrow), the *inter-burst interval* (red arrow), and *peak amplitude* (black arrow). *MEG*: magnetoencephalography; *ICA*: independent component analysis; *dSPM*: dynamic statistical parametric mapping; *ROI*: region of interest.

The ROI time-series was band-pass filtered between 13-30 Hz using a zero-phase finite impulse response filter to get the beta band time-course. The filter had a transition bandwidth of 3.25 Hz for the lower pass-band edge and a transition bandwidth of 7.5 Hz for the upper edge. We then applied a Hilbert transformation to the filtered time-series to obtain the instantaneous beta power.

### 2.5 Defining beta bursts

To asses and compare beta burst, we defined high-amplitude epochs in the envelope of the time-series above a fixed threshold defined in order of medians above the median of the envelope for each participant. To determine the value of the threshold, we took the correlation coefficient between the average amplitude of the signal envelope and the number of detected epochs within consecutive 3.0 seconds of data. This gave a single correlation coefficient per threshold per subject, which were averaged across all subjects. The threshold with the highest correlation between the number of epochs and signal amplitude was used as the fixed threshold in the comparisons (Fig. 1B). Defining the threshold in orders of medians, rather than an absolute cutoff value, gives a threshold that preserved the statistical properties at the group-level but fitted to the dynamic range of the individual subjects’ time-series. Similar methods for defining thresholds have been used to identify beta bursts in event-related studies (Feingold et al., 2015; Shin et al., 2017). Here we extended the method to resting-state MEG.

Once the threshold was defined, we extracted four features of the high-amplitude epochs (Fig. 1C). The first feature was the *rate* of occurrence within the three-minute time-series. The purpose of the first feature was to answer if the beta band were more “bursty” in one group compared to the other and whether it changed due to medication. The second feature was the epoch *duration*, defined as the time between the epoch reached the half-max of the peak value until it once again reached the half-max of the peak value (unless the half-max of the peak was above the threshold, in which case the time of threshold crossing was used to indicate the onset and offset). The purpose of the second feature was to answer if the high-amplitude epochs resembled “true” bursts (i.e., durations approximating one or two beta cycles) or perhaps showed prolonged high-amplitude activity in one of the groups. The third feature was the *inter-burst interval*, defined as the time from the offset of one epoch to the onset of the next even. The fourth and final feature was the *peak amplitude* of the envelope within each epoch.

### 2.6 Power spectral densities

To compare how the time-domain analysis compares to Fourier-based analysis of beta power in the frequency-domain, we calculated the PSD of the unfiltered ROI time-series in the spectrum from 1-48 Hz. We divided the time-series into consecutive epochs of three seconds with a 50% overlap and applied a Hanning taper before applying a fast Fourier transform using FieldTrip (Oostenveld et al., 2011) in MATLAB (R2016b; MathWorks Inc.).

### 2.7 Statistics

#### 2.7.1 Group characteristics

First, we tested for differences in age, sex ratio, and MoCA score between the Parkinson’s patients and healthy controls to ensure that the demographics of the two groups were adequately matched. Comparison of age and MoCA score by “Bayesian t-tests” (Rouder et al., 2009) using the *BayesFactor* package (Morey and Rouder, 2018) for R (R Core Team, 2013). The test gives the ratio of evidence for the hypothesis that there is a group difference versus the null-hypothesis of no difference between groups. To test for difference in the male-female ratio between groups, we used a Bayesian test for unequal multinomial distributions (Gûnel and Dickey, 1974).

#### 2.7.2 Power spectral densities

The PSDs were compared with pairwise cluster-based permutations tests across the spectrum from all 1-48 Hz. Independent t-test was first done on all frequency bins in the PSD. Adjacent t-values (df = 18 for within-group and df = 36 for between-group comparison) above or below the critical value (alpha < 0.05, two-tailed) were summed to gain the cluster T-value and then repeated on permutated datasets with randomized labels (n = 1000). The null hypothesis was rejected if the observed dataset had a largest cluster T-value above the 95th percentile of the permuted T-values (Maris and Oostenveld, 2007). The PSDs were compared across sessions within groups, between groups within sessions, and the interaction between groups and sessions. In addition to comparing the full spectrum between groups and sessions, we compared the relative power in the beta band by integrating the PSD in the beta range (13-30 Hz) and dividing it by integral of the full spectrum. The comparison of the relative beta power was done by pairwise Bayesian t-tests with the *BayesFactor* package in R.

#### 2.7.3 Beta burst features

The *rate*, *duration*, *inter-burst interval*, and *peak amplitude* were all analyzed by Bayesian mixed-effect regression, estimated in R with the *brms* package (Bürkner, 2017). The models used uninformative priors and were estimated by Markov-Chain Monte-Carlo sampling drawing 20.000 samples across four chains and discarding the first half of each chain. The convergence of the chains was confirmed by checking R̂ ≈ 1 (Gelman and Rubin, 1992).

We analyzed the epoch *rate* by mixed-effect Poisson regression containing Group (patient/control) and Session (first/second) as fixed effects with subjects as a random effect. The analysis of *duration*, *inter-burst interval*, and *peak amplitude* used the values for each epoch modeling the value of the *i*th epoch for participant *j* as a function of Group and Session by mixed-effect regression using the values of each epoch for all subjects. The *inter-burst interval* model used a log-normal link function, taking the log-transformed times to be Gaussian distributed. The models for *duration* and *peak amplitude* used shifted log-normal link functions that take the values subtracted a constant to follow a log-normal distribution.

Comparison between groups and sessions was done by comparing the marginal evidence—or Bayes factor (BF)—between models with and with the factor Group, Session, and the interaction between Group and Session as fixed effects. BF > 1 is evidence for the alternative hypothesis, whereas BF < 1 is evidence for the null-hypothesis. We use the guidelines by Wetzels et al., (2011) to determine the strength of the evidence where 0.33 < BF < 3 is taken as conclusive support for the alternative-or null-hypnosis. Values between 0.33 and 3 are inconclusive evidence. Post hoc hypothesis testing was done testing if at least 95% posterior distribution of individual parameters did not contain zero. The resulting test statistic is the probability *P* ranging from 0 to 1. *P* close to 0 is evidence for a difference between conditions, whereas *P* close to 1 provides evidence against a difference. We used the 95% posterior distribution corresponding to critical alpha = 0.05.

#### 2.7.4 Comparison across thresholds

To explore if the inference from the primary analysis was dependent on the threshold used to define the high-amplitude epochs, we repeated the comparison of the high-amplitude epoch rate between groups and sessions across thresholds. At each threshold—starting at the median to five times the order of median in steps of 0.1—we defined epochs as described above. The number of beta bursts at each threshold was analyzed by mixed-effect Poisson regression as in the primary analysis. We then compared models with and without the factor Group, Session, and the interaction between Group and Session to get a Bayes factor for each factor at each threshold. The model used uninformative priors and was estimated by Markov-Chain Monte-Carlo sampling drawing 4.000 samples across four chains and discarding the first half of each chain.

#### 2.7.5 Beta burst rate and motor symptoms

In addition to the group-level comparisons, we investigated the relationship between the burst rate and motor symptom severity measured with the MDS-UPDRS-III for the Parkinson’s patients. Since previous studies have shown that (frequency-domain) beta power is correlated with specific motor symptoms of rigidity and bradykinesia (Airaksinen et al., 2012, 2015; Melgari et al., 2014), we divided the MDS-UPDRS-III scores into six subscales of different motor symptoms according to the factors described by Goetz et al., (2008) with the exception that left-and right-side bradykinesia was combined into one factor. Each MDS-UPDRS-III factor (midline function, rest tremor, rigidity, bradykinesia, postural and kinetic tremor, lower limb bradykinesia) was modeled by mixed-effect Poisson regression as a linear function of the burst rate and subject and session as random intercepts. With these models, we tested the association between beta burst rate and the MDS-UPDRS-III factor scores by testing if at least 95% of the posterior distribution did not contain zero. All models model used uninformative priors and was estimated with *brms* (Bürkner, 2017) by Markov-Chain Monte-Carlo sampling drawing 20.000 samples across four chains and discarding the first half of each chain.

## 3 Results

### 3.1 Group characteristics

The groups are adequately matched for comparison as there were no systematic differences in the demographic variables: male/female ratio (BF = 0.60), age (BF = 0.41), and cognitive ability (BF = 0.39), see Table 1.

**Table 1:** Summary of the Parkinson’s group and control group.

|  | Parkinson's patients | Healthy controls |
| --- | --- | --- |
| *N* | 19 | 19 |
| *Sex* | 5 females, 14 males | 8 females, 11 males |
| *Age* | 44-85 years (mean: 67.3 years) | 54-76 years (mean: 69.3 years) |
| *Disease duration* | 1-14 years (median: 4.5 years) |  |
| *LEDD* | 300-1150 mg (median: 615 mg) |  |
| *MDS-UPDRS-III OFF* | 10-61 (median: 34) |  |
| *MDS-UPDRS-III ON* | 5-39 (median: 16) |  |
| *MoCA* | 25.5 (SD: 2.9) | (SD: 1.8) |

*LEDD*: levodopa equivalent daily dosage; *MDS-UPDRS-III*: Movement Disorder Society's Unified Parkinson's Disease Rating Scale part III; *MoCA*: Montreal Cognitive Assessment.

The Parkinson’s patients showed 26%-72% (mean 49%) improvement on motor symptoms on the MDS-UPDRS-III in the ON state compared to the OFF state (BF = 4.70*10^7^).

### 3.2 Power spectral densities

The cluster-based permutations test of the PSDs (Fig. 2) did not show any clusters of difference in any of the comparisons; thus, we cannot reject the null hypothesis that there is no difference between groups or sessions.

**Figure 2:**
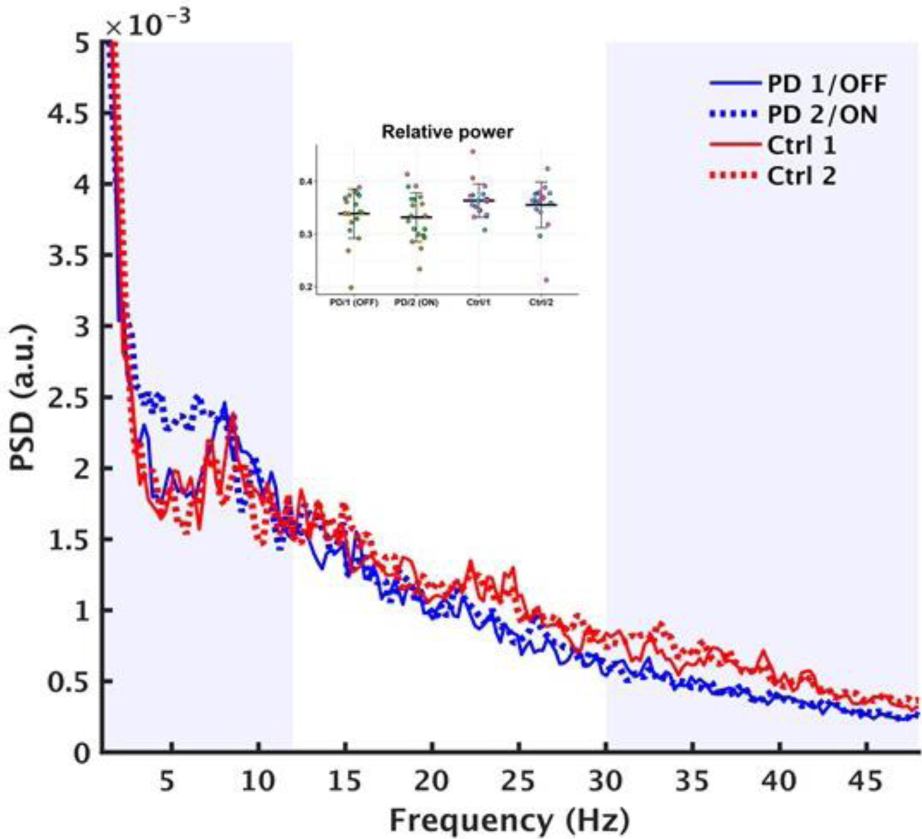
Group-level averaged power spectral densities. Parkinson’s patients in blue and healthy controls in red. solid lines is the first session/OFF medication and dashed lines is the second session/ON medication. The insert indicates the relative power of the beta band (13-30 Hz). *PSD*: power spectral density.

Comparison of the relative beta power gave evidence against a different between the first and second session for the controls (BF = 0.37) or between ON and OFF medication for the Parkinson’s patients (BF = 0.34). The comparison between the groups in the first session/OFF medication showed evidence for a difference between the groups but only as inconclusive evidence (BF = 1.27) and gave inconclusive evidence against a difference between the groups in the second session (BF = 0.87). Based on the comparisons in the frequency domain, we are not able to conclude that there is a difference between groups or between sessions.

### 3.3 Beta burst rate

The Parkinson’s patients showed an average rate of 106 bursts/min (SD: 8) in the first session/OFF medication and 108 bursts/min (SD: 11) in the second session/ON medication. The controls had an average rate of 120 bursts/min (SD: 11) in the first session and 116 bursts/min (SD: 15) in the second session. Fig. 3 shows the burst rate for all subjects across groups and sessions.

**Figure 3:**
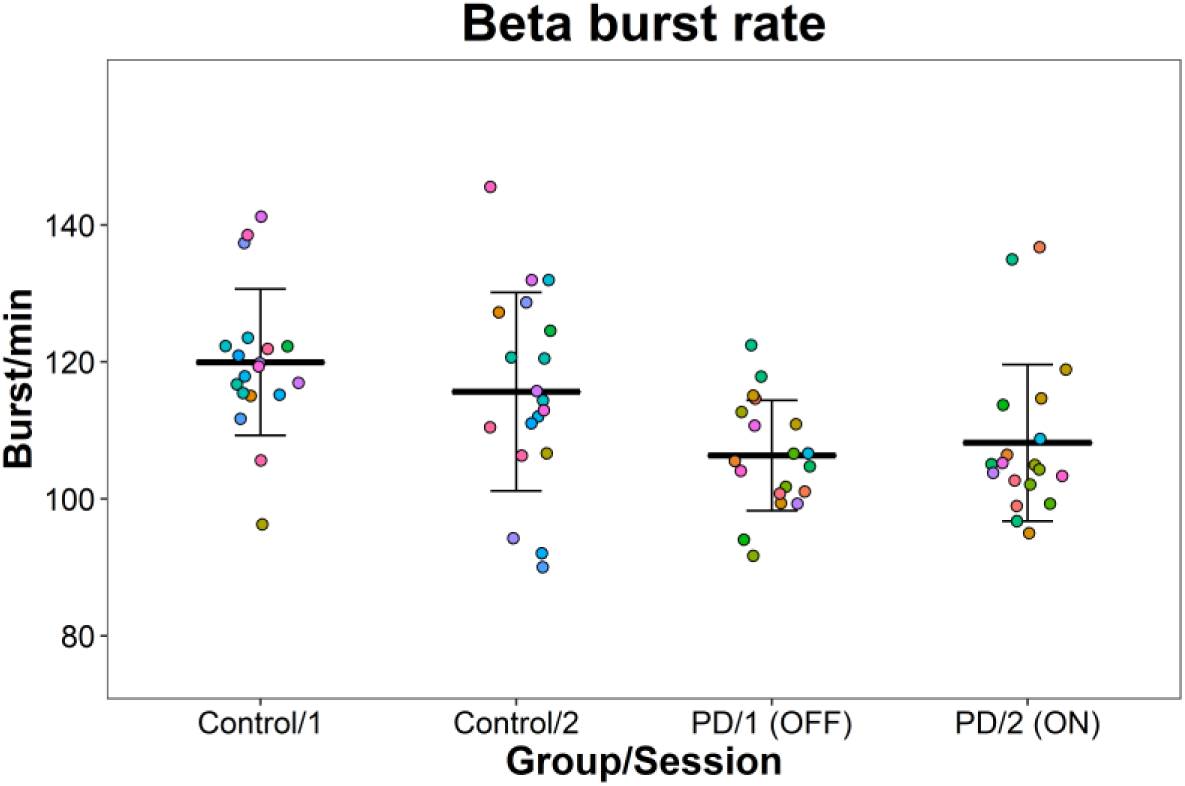
Beta burst rate in the sensorimotor areas across groups and sessions. The points represent the beta bursts rate for each participant. The bars are means and standard deviations.

The model comparison showed evidence for an effect of Group (BF = 10.9) but gave evidence against an effect of Session (BF = 0.062) and gave evidence against interaction between Group and Session (BF = 0.24).

The Parkinson’s patients had 5-17% (median: 11%) lower rate in the OFF state compared to healthy controls (P < 0.0016). The change in rate from the OFF to the ON state varied from a 4% reduction to 8% increase (median 2% increase) and was not significantly different from zero (P = 0.60). The healthy controls showed a change in burst rate from the first to the second that ranged from a 9% decrease to a 2% increase (median: 3% decrease). The change in burst rate between session for the healthy controls was not significantly different from zero (P = 0.22).

### 3.4 Burst duration

The high-amplitude epochs showed that the beta bursts were short, with a median duration between 73-76 ms in both sessions and groups (see Table 2). 95% of the epoch duration distributions fell within 35-170 ms. The median duration of the beta bursts corresponded roughly to a single oscillatory cycle in the beta frequency range (approximately 13-14 Hz).

**Table 2:** Group-level summary of beta burst features (medians and 95%-predictive intervals).

| Group-session | Bursts/min | Duration | Inter-burst interval | dSPM peak amplitude |
| --- | --- | --- | --- | --- |
| *Parkinson's patients 1/OFF* | 106 (86-128) | 74 ms (36-165) | 184 ms (9-3903) | 0.99 (0.61-1.74) |
| *Parkinson's patients 2/ON* | 108 (88-130) | 76 ms (37-170) | 177 ms (9-3686) | 1.00 (0.61-1.77) |
| *Healthy controls 1* | 120 (98-142) | 73 ms (36-163) | 136 ms (7-2887) | 0.95 (0.59-1.65) |
| *Healthy controls 2* | 116 (95-138) | 73 ms (35-159) | 147 ms (7-2901) | 0.98 (0.60-1.73) |

*dSPM*: dynamic statistical parametric map

The comparison of the burst durations showed evidence against an effect of Session (BF = 0.046), gave evidence against an effect of Group (BF = 0.17), and gave evidence against the interaction between Session and Group, though the evidence is in the inconclusive range (BF = 0.59).

### 3.5 Inter-burst intervals

The inter-burst intervals had a skewed distribution with a high probability of short intervals below 200 ms with few longer intervals that could last up to seconds (Fig. 4B). The model comparison showed evidence against an effect of Session (BF = 0.049) and evidence for an effect of Group (BF = 283). For the inter-burst intervals, there was evidence for an interaction between Group and Session (BF = 5173).

**Figure 4:**
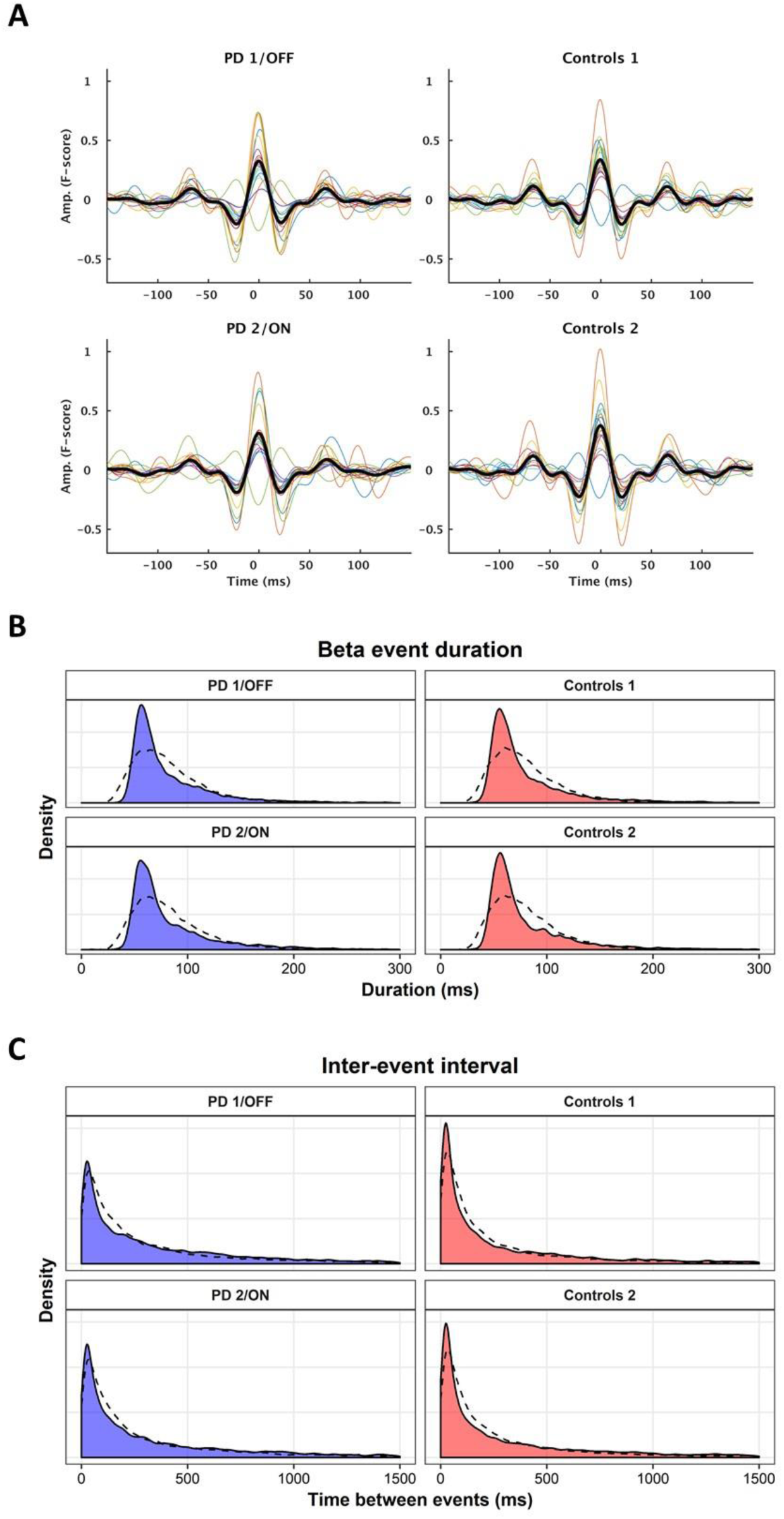
Beta burst features. A) Average beta bursts time-locked to the burst peak for each group/session. Thick lines are the grand average, and colored lines are individual subjects. Pooled distributions of the burst duration (B) and inter-burst intervals (C) across groups and sessions. Dashed lines in (B) and (C) are the group-level predicted values of the models.

The model showed a median inter-burst interval of 187 ms (mean: 631 ms, 95%-I: 9-3903 ms) for patients OFF medication, compared to a median inter-burst interval of 136 ms (mean: 442 ms, 95%-I: 7-2887 ms) for healthy controls in the first session (P < 10^-4^). The median inter-burst interval decreased to 177 ms (mean: 560 ms, 95%-I: 9-3686 ms) in the ON medicated, corresponding to a 10% decrease (CI: 4%-14%) in the inter-burst intervals from the OFF to ON medication state (P = 2*10^-4^). The inter-burst interval changed in the opposite for the healthy controls and increased by 8% (CI: 3-14%) between sessions (P = 0.003).

### 3.6 Peak amplitude

Fig. 4A depicts averaged beta burst time-locked to the peak amplitude. The peak amplitude of the beta bursts only differed between sessions independent of the group. The model comparison of the peak amplitude showed evidence for an effect of Session (BF = 1.6*10^9^), but evidence against an effect of Group (BF = 0.46) and evidence against the model that included the interaction between Session and Group (BF = 0.48)—though the BFs are in the inconclusive range for the two latter model comparisons. The peak amplitude increased for both controls and patients in the second session with an increase of 4% (CI: 3-5%; P < 10^-4^) for controls and an increase of 2% (CI: 1-3%; P = 0.002) for the Parkinson’s patients.

### 3.7 Comparison across thresholds

To investigate how the threshold for defining beta bursts influenced the inference, we repeated the comparison of the burst rate across a range of thresholds. Fig. 5B shown the Bayes factors of the comparison across the thresholds. The model comparisons for all thresholds above one unit of medians favored a difference in the number of beta bursts between controls and patients with the patients having fewer beta bursts than the controls. At higher thresholds, the comparison favored and interaction between Group and Session, with an increase in the burst rate from OFF to ON but also increased variation (Fig. 5A). Since the inference one would draw at different thresholds is consistent across thresholds (with the exception of the very low and high thresholds), we conclude that the inference is not too dependent on the precise numerical threshold.

**Figure 5:**
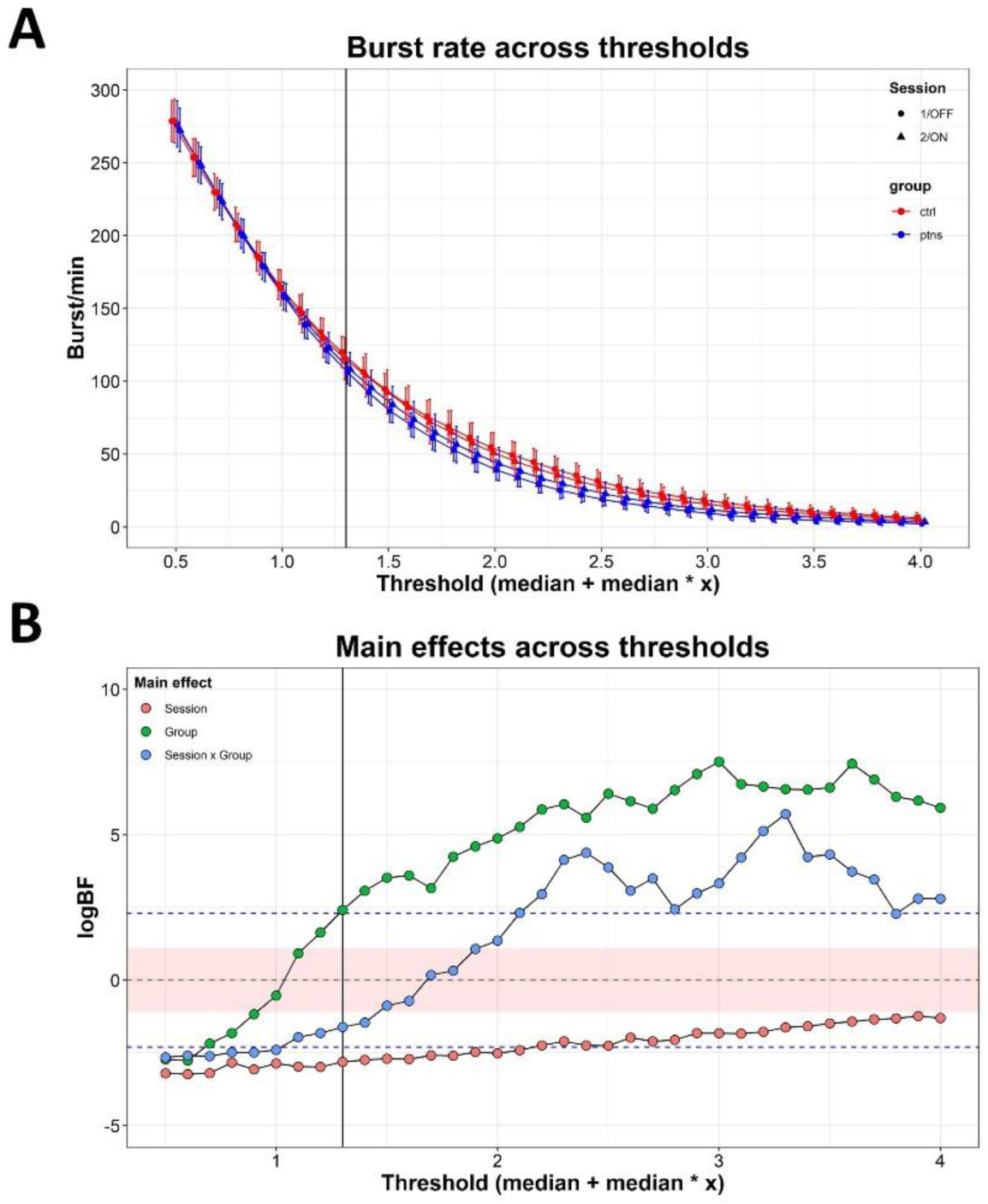
Comparison across thresholds for defining beta bursts. A) The beta burst rate depending on the thresholds used to define beta bursts for both groups and sessions. B) The results of the Bayesian-model comparison across thresholds. The red area indicates the interval where the Bayes factors are considered inconclusive for or against the hypothesis and the dashed red lines indicate “substantial” evidence for (upper line) or against (lower line) the hypothesis, following the guidelines by Wetzels et al. (2011). The vertical line indicates the threshold used in the primary analysis. *logBF*: logarithm of Bayes factor.

### 3.8 Beta burst rate and motor symptoms

Fig. 6 shows the marginal predicted effects of the burst rate and the subscales of the MSD-UPDRS-III from the regression models. The burst rate scaled negatively with bradykinesia (P = 0.038). The regression model predicted a decrease of 29% (95%CI: 10-45%) in bradykinesia rating when the burst rate increased by 10. The burst rate further scaled negatively with postural/kinetic tremor (P = 0.028), predicting 40% (95%CI: 16-59%) decrease in symptom rating when the burst rate increased by 10. We saw no evidence that midline function (P = 0.44), rest tremor (P = 0.71), rigidity (P = 0.87), nor lower limb bradykinesia (P = 0.28) scaled with the burst rate.

**Figure 6:**
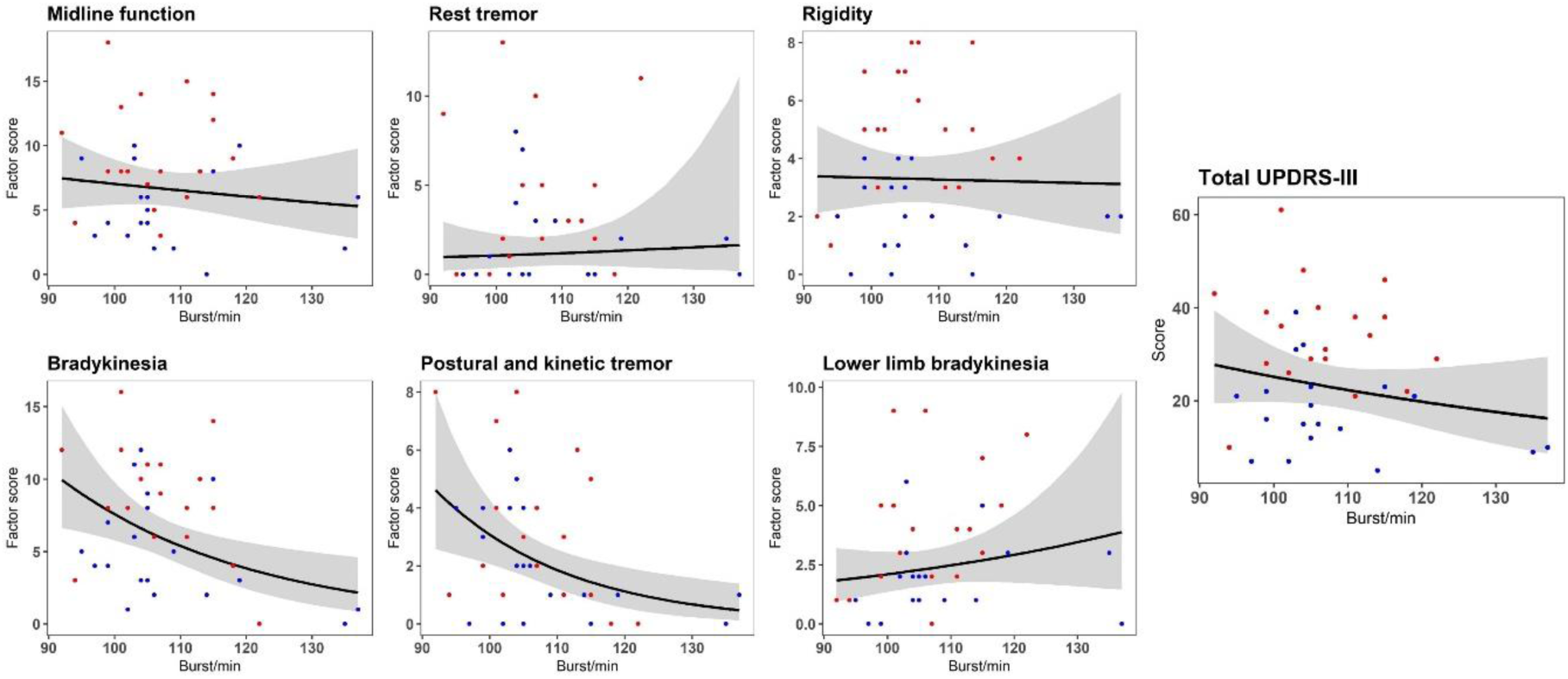
Relation between the beta bursts rate and MDS-UPDRS-III subscales. Colored dots are individual measurements OFF (red) and ON (blue) medication. Lines and shades indicate the predicted marginal effect of burst rate on the score on MDS-UPDRS-III subscales. *MDS-UPDRS-III*: Movement Disorder Society’s Unified Parkinson’s Disease Rating Scale, part III.

## 4 Discussion

The primary aim of this study was to explore whether beta burst characteristics differed between Parkinson’s patients and healthy controls. As a secondary aim, we also explored whether beta burst characteristics vary within Parkinson’s patients because of dopaminergic medication; and finally, as a third aim, explored whether beta burst characteristics were related to symptom severity in Parkinson’s disease.

When the Parkinson’s patients were OFF medication showed a 6-17% lower beta burst rate compared to healthy controls. This reduction of in beta burst rate was still present when the patients were ON medication. Neither the duration nor the amplitude of the beta bursts differed between patients and controls. Our results add to the evidence that the cortical activity in the beta band exhibits transient bursts lasting a one or two cycles. This is in line with the research from Sherman et al. (2016), who proposed that beta burst in the cortex is caused by a short distal drive in the upper laminar layers lasting around 50 ms in combination with a sustained excitatory proximal drive between the upper and lower cortical layers. The consistency in duration and amplitude suggests that some components of the mechanisms that generate the cortical beta bursts are preserved in Parkinson’s disease, while the rate of bursts decreases with the disease and with symptom severity. This reduction in spontaneous beta bursts in the sensorimotor cortex could potentially be driven by a reduction in distal connections from thalamus or the basal ganglia.

The distribution of the inter-bursts interval more resembled the distributions of the healthy controls when patients were on medication. What this means in terms of disease-related mechanisms is currently unclear as the underlying dynamics that drive the beta bursts is unknown. It is possible that the shift in inter-burst interval following dopaminergic medication is driven by a change in the distal drive from dopamine modulated activity in basal ganglia or thalamus. However, more research is needed to understand how the cortical beta bursts are driven by deeper sources, which directions the connection goes, and how this is modulated by dopaminergic medication. The effect of dopaminergic medication on the beta band seems to be much more complex than changes in the average power in the beta band. Rather, by assessing the beta activity in terms of beta burst and analyzing the characteristics of these events, it seems that what mainly changes in the temporal distribution of transient beta bursts.

We did not find an effect of dopaminergic medication on the burst rate, nor on the burst duration. Since the study was exploratory and we did not have prior estimates of an expected effect size of medication and that our sample size was relatively small (n = 19), there might be effects of medication that we have not detected with this analysis approach. At higher thresholds than the one used in the main analysis (Fig. 5), there was evidence for an effect of medication on the burst rate.

In the Parkinson’s patients, the decrease in beta burst rate was associated with an increase in symptom severity for bradykinesia and postural/kinetic tremor. Such a link between burst rate and bradykinesia is in line with previous studies showing that decreased beta power in the cortex is related to increased bradykinesia (Airaksinen et al., 2012, 2015; Melgari et al., 2014). A reduction in the average PSD is compatible with the reduction in the number of spontaneous high-amplitude bursts as well as a reduction in sustained oscillatory activity. However, in our results, we did not observe any conclusive differences between Parkinson’s patients and healthy controls in the averaged PSD that corresponds to those we report for the analysis of beta bursts (only an inconclusive trend for the relative beta power). For our data, frequency-domain analysis using the traditional Fourier-transform method thus seems to be less sensitive in picking up statistically meaningful differences in beta activity between Parkinson’s patients and healthy controls compared to an analysis based on individual burst events.

The variation between the sessions for the healthy controls may reflect the test-retest variability of the measurements, which were between a 9% decrease to a 2% increase in beta burst rate. This variation can also reflect a circadian effect on the spontaneous beta bursts. It has previously been shown that the frequency domain beta power varies with the time of the day (Wilson et al., 2014). It is plausible that similar circadian effects apply to beta bursts in the time-domain. All participants—Parkinson’s patients and controls alike—were tested in the morning and again before noon on the same day in our study.

The presence of cortical beta band activity is inversely related to motor function: a decrease in beta band activity indicates an increased sensitivity to efferent and afferent sensorimotor signals, whereas increased activity has been linked to inhibition of sensorimotor signals (Brown, 2007; Engel and Fries, 2010). Close temporal proximity between beta bursts and go cues leads to longer reaction times (Little et al., 2018; Lofredi et al., 2019) and less likelihood of detecting sensory stimuli close to the sensory threshold (Shin et al., 2017), suggesting that the proximity of beta bursts blocks immediate sensorimotor processing. Spontaneous beta bursts thus seems to have a transient inhibitory effect on the sensorimotor processing, but might at the same time serve as a signal that is necessary to maintaining a continuous optimal state of sensorimotor processing (Engel and Fries, 2010; Jenkinson and Brown, 2011). This interpretation entails that the beta bursts serve as an immediate updating of the sensorimotor system by integrating the previous motor signal and proprioceptive signal (Leventhal et al., 2012). The beta bursts might be inhibitive, as evidenced by their behavioral effects on event-related sensorimotor tasks (Shin et al., 2017; Little et al., 2018), but keeping maintenance of the sensorimotor system over a longer time. The inverse relation between the number of spontaneous beta bursts and bradykinesia, that we report in this study, might hence be due to a deficit in the updating of the sensorimotor system, which leads to suboptimal utilization of neural resources when initiating and performing movements manifesting as bradykinesia and kinetic tremors.

It is well known that beta activity is altered in Parkinson’s disease, which is often evident at the frequency domain on decomposed and averaged time-series of electrophysiological activity. However, that approach implicitly assumes that the average signal is representative of the whole time-series. The neuronal oscillations in the beta band change over time by exhibiting transient beta bursts lasting 70-80 ms. We have shown that the burst duration is similar for both healthy adults and Parkinson’s patients—but that the *burst rate* is reduced in Parkinson’s disease. The spontaneous dynamics in the beta band, such as burst events and burst characteristics, might hold further information that is relevant for understanding Parkinson’s disease and the development of motor symptoms. Modulation of the dynamic changes in the beta activity due to the dopaminergic medication has been shown in deep-brain recordings from STN in Parkinson’s patients (Tinkhauser et al., 2017a, b). Recordings of the electrical field in STN is only done in patients who undergo brain surgery. It is, therefore, not feasible for diagnostic purposes. Here we show that Parkinson’s patients exhibit a reduction in the beta bursts rate compared to healthy controls and that this can be measured from the cortex non-invasively using MEG.

## Acknowledgments

Data for this study was collected at NatMEG, the National Facility for Magnetoencephalography (http://natmeg.se), Karolinska Institutet, Sweden. The NatMEG facility is supported by Knut & Alice Wallenberg (grant #KAW2011.0207). We are thankful to Allison Eriksson for proofreading the manuscript.

## Data Availability

The datasets collected for the current study contains patient information that cannot be made public. The dataset is available from the corresponding author for review purpose or on reasonable request. Scripts for running the analysis presented in the paper is available at www.github.com/mcvinding/PD_beta_bursts.

